# Generation and application of river network analogues for use in ecology and evolution

**DOI:** 10.1101/2020.02.17.948851

**Authors:** Luca Carraro, Enrico Bertuzzo, Emanuel A. Fronhofer, Reinhard Furrer, Isabelle Gounand, Andrea Rinaldo, Florian Altermatt

## Abstract

1. Several key processes in freshwater ecology and evolution are governed by the connectivity inherent to dendritic river networks. These networks have extensively been analyzed from a geomorphological and hydrological viewpoint, yet network structures classically used in modelling have only been partially representative of the structure of real river basins, and have often failed to capture well known scaling features of real river networks. Pioneering work has identified optimal channel networks (OCNs) as spanning trees that reproduce all scaling features characteristic of real, natural stream networks worldwide. While these networks have been used to generate landscapes for studies on metapopulations, biodiversity and epidemiology, their generation has not been generally accessible.
2. Given the increasing interest in dendritic riverine networks by ecologists and evolutionary biologists, we here present a method to generate OCNs and, to facilitate its application, we also provide the R-package OCNet. Owing to the random search process that generates OCNs, multiple network replicas spanning the same surface can be built, allowing one to perform computational experiments whose results do not depend on the particular shape of a single river network. The OCN construct also enables the generation of elevational gradients derived from the optimal network configuration, which can constitute three-dimensional landscapes for spatial studies in both terrestrial and freshwater realms. Moreover, the OCNet package provides functions that aggregate the OCN into an arbitrary number of nodes, calculate several metrics and descriptors of river networks, and draw relevant features of the network.
3. We describe the main functionalities of the package and present how it can be integrated into other R-packages commonly used in spatial ecology. Moreover, we exemplify the generation of OCNs and discuss an application to a metapopulation model for an invasive riverine species.
4. In conclusion, OCNet provides a powerful tool to generate and use realistic river network analogues for various applications. It thereby allows the design of spatially realistic studies in increasingly impacted ecosystems, and enhances our knowledge on spatial processes in freshwater ecology in general.

## 1 Introduction

The central goal of ecology is to causally understand patterns and processes in ecological systems, such as species coexistence, biodiversity patterns, or the unfolding of species invasions [Gause, 1934; Elton, 1958]. Much of ecological theory and empirical work has either focused on local patterns and dynamics or has taken a spatially implicit perspective. However, virtually all natural ecosystems are spatially structured, and the relevance of the spatial dimension on ecological systems can hardly be overestimated [Levin, 1992; Hanski and Gaggiotti, 2004; Holyoak et al., 2005]. Consequently, over the last decades, ecologists have started to account for spatial processes on population and community dynamics as well as biodiversity. Theoretical, comparative and experimental studies have increasingly been done in a spatially explicit perspective (e.g., Hanski and Ovaskainen [2000]; Cadotte and Fukami [2005]; Holyoak et al. [2005]; Altermatt et al. [2011]; Gilarranz and Bascompte [2012]; Dale and Fortin [2014]), especially promoted by theories on metapopulation and metacommunity dynamics.

A direct consequence of this spatial approach to ecology is the need to describe and understand the spatial structure and layout of natural ecosystems. While initial models of spatial dynamics assumed spatially implicit networks of populations or communities [Levins, 1970], all natural ecosystems follow spatially explicit structures. These structures, such as those typically found in coral reefs and atolls, mountainous landscapes and their elevational gradients, or tidal pools, are shaped by general geophysical processes resulting in characteristic landscape structures. Arguably the most characteristic (but also among the most widespread) landscape structure is found in riverine networks [Leopold et al., 1964; Rodriguez-Iturbe and Rinaldo, 2001]: erosional forces balancing uplift create dendritic networks of rivers and streams following universal patterns. These networks are characterized by their fractal, scale-free structure, as well as by universally applicable laws regarding many geomorphological and hydrological variables of direct relevance to ecology, such as catchment area, river bed width and depth, or variation in discharge [Horton, 1945; Leopold and Maddock, 1953; Rodriguez-Iturbe and Rinaldo, 2001]. In contrast to these specific features of natural landscape structures, much of ecological and evolutionary theory and experiments, but also much of the species-distribution modelling still assumes either random networks or simply structured linear, circular or Cartesian networks, in which local patches are connected to their 2, 4 or 8 nearest neighbours [e.g. Bascompte and Solé, 1996; Bell and Gonzalez, 2011; Holland and Hastings, 2008]. This oversimplification of spatial network structures may limit the plausibility and relevance of the findings. An application to more realistic network structures has, however, often been hindered by the lack of formalized, spatially correct and generalizable network structures as well as easily accessible tools generating them.

Riverine ecosystems are not only of high interest to ecologists due to their universal network structure, but also due to the considerable biodiversity inhabiting them [Balian et al., 2008; Altermatt, 2013]. River networks cover less than 1% of the landmasses, but contain up to 10% of all species. However, this high biodiversity, as well as the associated ecosystem functions, are threatened by various anthropogenically induced causes, including pollution, biological invasions, or damming and modification of the network structure [Vörösmarty et al., 2010; Darwall et al., 2018]. An understanding of many of these processes requires a spatially explicit approach, such as how pollution and chemicals are transported in riverine networks [Helton et al., 2018], how organisms spread along rivers and invade riverine ecosystems [Mari et al., 2014; Giometto et al., 2017], or how the modification of network structures across drainage basins affects local diversity [Leuven et al., 2009]. Consequently, there has been a rapid increase in ecological and evolutionary studies considering the effect of river-like network structures on ecological dynamics over the last two decades [Fagan, 2002; Campbell Grant et al., 2007], paralleled by an increase in methodological tools to analyse such spatial datasets [Muneepeerakul et al., 2008; Rodriguez-Iturbe et al., 2009; Peterson et al., 2013; Paz-Vinas and Blanchet, 2015; Welty et al., 2015; Duarte et al., 2019; Rinaldo et al., 2020].

While all of these works acknowledge the importance of studying rivers in a spatially explicit perspective, a large part of them is built on networks that do not factually capture many of the inherent characteristics of true riverine networks. Notable examples range from the River Continuum Concept [Vannote et al., 1980], which describes rivers as a single, linear array of patches, to slightly more complex bifurcation networks or alterations thereof [Fagan, 2002; Morrissey and De Kerckhove, 2009; Chaput-Bardy et al., 2009; Brown and Swan, 2010; Yeakel et al., 2014; Seymour and Altermatt, 2014; Seymour et al., 2015; Paz-Vinas et al., 2015; Anderson and Hayes, 2018]. All of these studies use networks that may at first sight look like “river” networks, but do not satisfy the necessary constraint posed by draining a given surface (Fig. 1). Furthermore, these constructs do not adequately represent the connectivity and several geometric properties (like the distributions of upstream and downstream lengths, and of total contributing area at a point) inherent to natural river networks, and lack the space-filling attribute of small to smallest streams not only incrementally flowing into larger streams, but also the common direct inflow of very small streams into large streams. As such, all of this work has been ignoring the extensive and long-lasting knowledge from geomorphology that has appropriately acknowledged and formalized the spatial unfolding of dendritic river networks.

**Figure 1:**
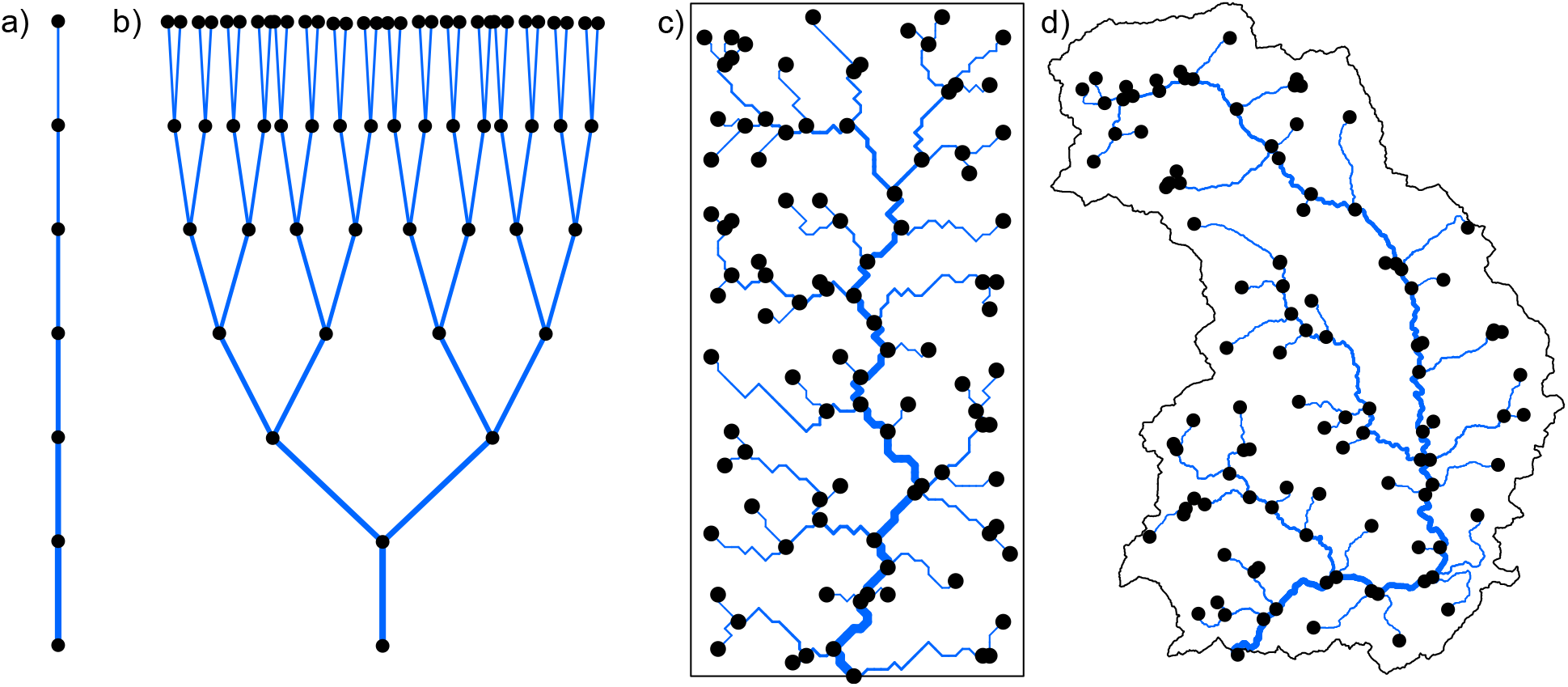
Examples of river network analogues with increasing level of resemblance with real river networks. Line width increases towards the downstream direction. a) Linear array of patches [Vannote et al., 1980]. b) Binary-fission-like tree [Fagan, 2002; Paz-Vinas et al., 2015]. c) An OCN spanning a 50×100 lattice, aggregated with a threshold area equal to 25 pixels. d) A real river (Thur, Switzerland), spanning an area of 760 km^2^, extracted from a digital elevation model with a threshold area equal to 4 km^2^.

In particular, the fractal character of river networks, epitomized by Horton’s laws [Horton, 1945] on bifurcation and length ratios, was observed with regards to several morphological and hydrological characteristics of river basins and expressed by means of a number of power-law relationships, which are the signatures of fractal behaviour [Mandelbrot, 1983; Maritan et al., 1996]. Notable examples are Hack’s law [Hack, 1957] *L* ∼ *A*^*h*^, linking the maximum upstream channelized length *L* at any location in the river with the corresponding drainage area *A*; the slope-area relationship *s* ∼ *A^γ^*^−1^, where *s* is the channel slope [Tarboton et al., 1989]; the scaling of the probability distribution of drainage areas *P*(*A* ≥ *a*) ∼ *a*^−*β*^ [Rodriguez-Iturbe et al., 1992a]. Typical values observed in real rivers for the scaling exponents are *h* ≈ 0.57, *γ* ≈ 0.5, *β* ≈ 0.43 [Rinaldo et al., 2014]. From a hydraulic geometry viewpoint, Leopold’s relationships [Leopold and Maddock, 1953] express how mean river depth, width and velocity change, both at-a-station and along the river’s course, as power-law functions of discharge, which in turn scales linearly with drainage area (strictly speaking, this applies to landscape-forming discharges [Rodriguez-Iturbe and Rinaldo, 2001]).

Such scale-invariant properties of river networks prompted the development of a model of idealized stream networks: optimal channel networks (OCNs). OCNs are ‘optimal’ inasmuch as their configuration corresponds to a minimum of total energy expenditure and reproduces all scaling features of real rivers [Rodriguez-Iturbe et al., 1992b; Rinaldo et al., 1992; Maritan et al., 1996; Rinaldo et al., 2014]. Importantly, OCNs are exact stationary solutions of the general equation describing landscape evolution [Banavar et al., 2001]. The OCN construct allows the generation of an unlimited number of different network replicas spanning the same drainage domain, therefore enabling one to run computational experiments and derive results that are independent of the shape of a single river network, which would not be the case if real rivers were used as landscapes. Moreover, OCNs enable the investigation of spatial processes occurring not only in dendritic river networks, but also along the elevational gradients of fluvial landscapes [Bertuzzo et al., 2016; Giezendanner et al., 2019]. To this regard, it is worthwhile to note that the elevational landscape generated by an OCN is such that the graph obtained by following the steepest descent directions reproduces the OCN structure [Balister et al., 2018].

OCNs have been used to investigate a number of ecological issues, ranging from metapopulation structure in riverine [Mari et al., 2014; Bertuzzo et al., 2015; Fronhofer and Altermatt, 2017] and terrestrial landscapes [Bertuzzo et al., 2016; Giezendanner et al., 2019]; habitat fragmentation [Sarker et al., 2019]; spreading of human [Bertuzzo et al., 2010; Gatto et al., 2013; Mari et al., 2019] and animal [Carraro et al., 2018] waterborne pathogens; ecosystem processes, such as carbon [Bertuzzo et al., 2017; Koenig et al., 2019] and nitrogen cycling [Helton et al., 2018]; migration fronts of human populations [Campos et al., 2006]; cross-ecosystem subsidies [Harvey et al., 2019]; riverine biodiversity patterns from a theoretical viewpoint [Muneepeerakul et al., 2019] or by means of mesocosm experiments [Carrara et al., 2012, 2014; Harvey et al., 2018].

Despite the longstanding establishment of the OCN concept, its application especially in ecology and evolutionary biology has been lagging behind, likely because easily accessible code or appropriate tools have been lacking. This is particularly regrettable considering the recent bloom of tools allowing the statistical analysis of data from real dendritic networks (e.g. the R-package SSN [Ver Hoef et al., 2014]). However, such tools are specifically designed for real river networks, while their applicability to virtually generated networks is limited. To fill this gap, we here describe the methodological and mathematical frameworks that underlies `OCNet`, an R-package for the generation and analysis of optimal channel networks, and provide guidelines and examples to facilitate the use of this tool.

## 2 The OCNet package

### 2.1 Theoretical background

The OCN concept is based on the assumption that river network configurations occurring in nature minimize a functional describing total energy dissipation [Rodriguez-Iturbe et al., 1992b; Rinaldo et al., 1992]. This assumption, and the subsequent construction of OCNs, is well supported by a comparison with river networks globally. Let us consider a regular lattice made up of *N* cells, where each cell represents the generic node *i* of the network. Each node *i* is connected via a link to one of its nearest neighbours. The energy dissipation across the *i*th network link is proportional to *Q*_*i*_Δ*h*_*i*_, where *Q*_*i*_ is the landscape-forming discharge in the link [Rinaldo et al., 2014], and Δ*h*_*i*_ = *s*_*i*_L_i_ the corresponding elevation drop, with *s*_*i*_ identifying slope and *L_i_* link length. By assuming *Q*_*i*_ ∼ *A_i_* [Rodriguez-Iturbe and Rinaldo, 2001], where *A*_*i*_ is the area drained by link *i*, and the slope-area relationship 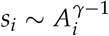 [Tarboton et al., 1989], the functional representing total energy expenditure across a landscape formed by *N* cells reads

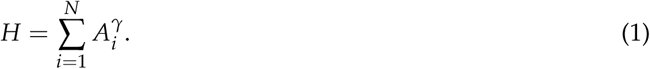

Note that link lengths do not appear in the above formula, as they can be considered constant with no loss of generality. The OCN configuration is defined by an adjacency matrix **W** whose corresponding set of drainage areas **A** = [*A*_1_, …, *A*_*N*_] yields a local, dynamically accessible minimum of (1). Note that the correspondence between **A** and the adjacency matrix **W** of a tree is subsumed by the relationship (**I**_*N*_ − **W**^*T*^)**A** = **1**, where **I**_*N*_ is the identity matrix of order *N*, and **1** a *N*-dimensional vector of ones [Bertuzzo et al., 2015].

Minimization of (1) is operated by means of a simulated annealing technique: starting from a feasible initial flow configuration (i.e., a spanning tree, see Fig. 2a, e), a link at a time is rewired to one of its nearest neighbours; if the obtained configuration is a spanning tree, *H* is computed; the new configuration is accepted if it lowers total energy expenditure; if this is not the case, the new configuration can still be accepted with a probability controlled by the cooling schedule of the simulated annealing algorithm. Such myopic search, which only explores close configurations, actually mimics the type of optimization that nature performs, at least in fluvial landscapes [Rinaldo et al., 2014]. Notably, restricting the search of a network yielding a minimum of Eq. (1) to spanning, loopless configurations entails no approximation, because every spanning tree is a local minimum of total energy dissipation [Banavar et al., 2000]. The shape of the so-obtained OCNs retains the heritage of the initial flow configuration, although the extent to which this occurs is partly controlled by the cooling schedule adopted (Fig. 2). This underpins the concept of feasible optimality, i.e., the search for dynamically accessible configurations.

**Figure 2:**
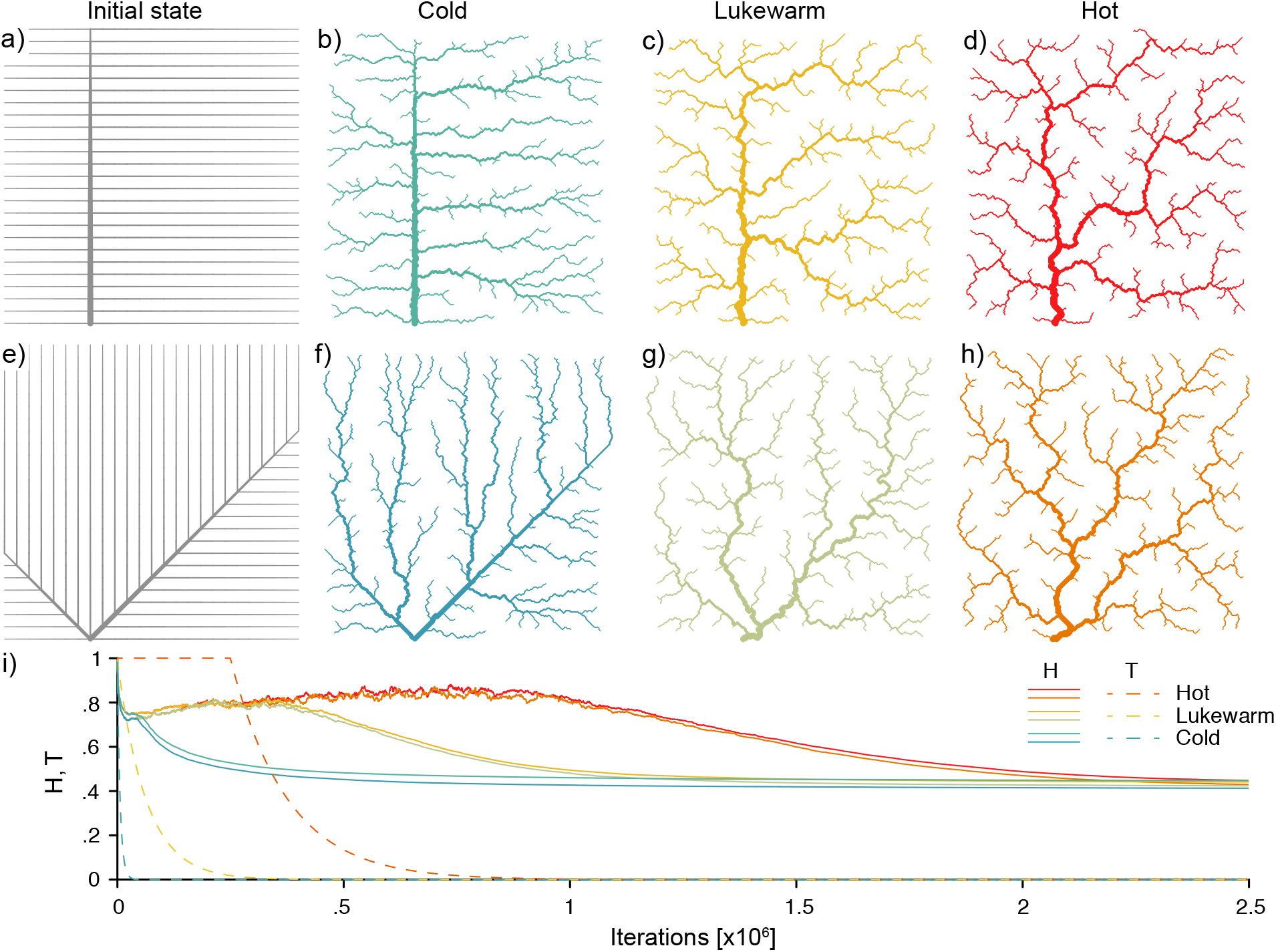
Effect of initial network state (rows) and cooling schedule (columns) on the final OCN configuration. b, c, d) OCNs on 250×250 lattices generated from the initial state shown in panel a. f, g, h) As above but with initial state as shown in panel e. i) Dynamics of total energy expenditure *H* (Eq. (1)) and temperature *T* (i.e., cooling schedule of the simulated annealing algorithm) for the 6 OCNs displayed above. Values of *H* and *T* are normalized by the energy of the initial network state. Note that, for graphical reasons, the initial states shown in panels a, e refer to 25×25 lattices. A script generating this figure can be found in the Supporting Information.

### 2.2 Overall setup of the package

The `OCNet` package consists of a series of functions that allow constructing river-analogue networks as well as calculating a number of metrics and descriptors commonly used in spatial ecology. The networks constructed by the package are built at several levels of aggregation. At each level, they are generally defined by a number of nodes, an adjacency matrix, a vector of contributing areas and two vectors with longitudinal and latitudinal coordinates of the nodes. The functions constituting the `OCNet` package are intended to be applied in sequential order, and the respective output can be directly used to visualize the created networks and linked to other commonly used R-packages.

The first function, `create_OCN`, only requires the longitudinal and latitudinal dimensions of the lattice as necessary inputs, while several other parameters can be optionally tuned to obtain customized results. Some examples are provided in the following section; extensive further information is given in the package documentation. The output of `create_OCN` is a list containing a sub-list termed FD that, in turn, encloses key information on the topology of the network, among which the adjacency matrix (written in sparse form via the `spam` format [Furrer and Sain, 2010]) and a vector of contributing areas. The subsequent functions `landscape_OCN` (generation of the three-dimensional landscape derived from the network configuration), `aggregate_OCN` (aggregation of the OCN at various levels – see *Aggregation levels*), paths OCN (evaluation of paths among network nodes, and lengths thereof), `rivergeometry_OCN` (hydraulic geometry of the river network, following Leopold and Maddock [1953]) require as necessary input the output list produced by the previous function, in the aforementioned order (except `rivergeometry_OCN`, which can be executed after `aggregate_OCN`). The output of these functions is a list where all objects of the input list are copied, and to which new objects are added. Note that output lists contain all input values, to avoid inconsistencies in the sequential application of functions. A group of functions (identified by the prefix `“draw_”`, see examples in Fig. 3) are devoted to graphical representations of the OCN.

**Figure 3:**
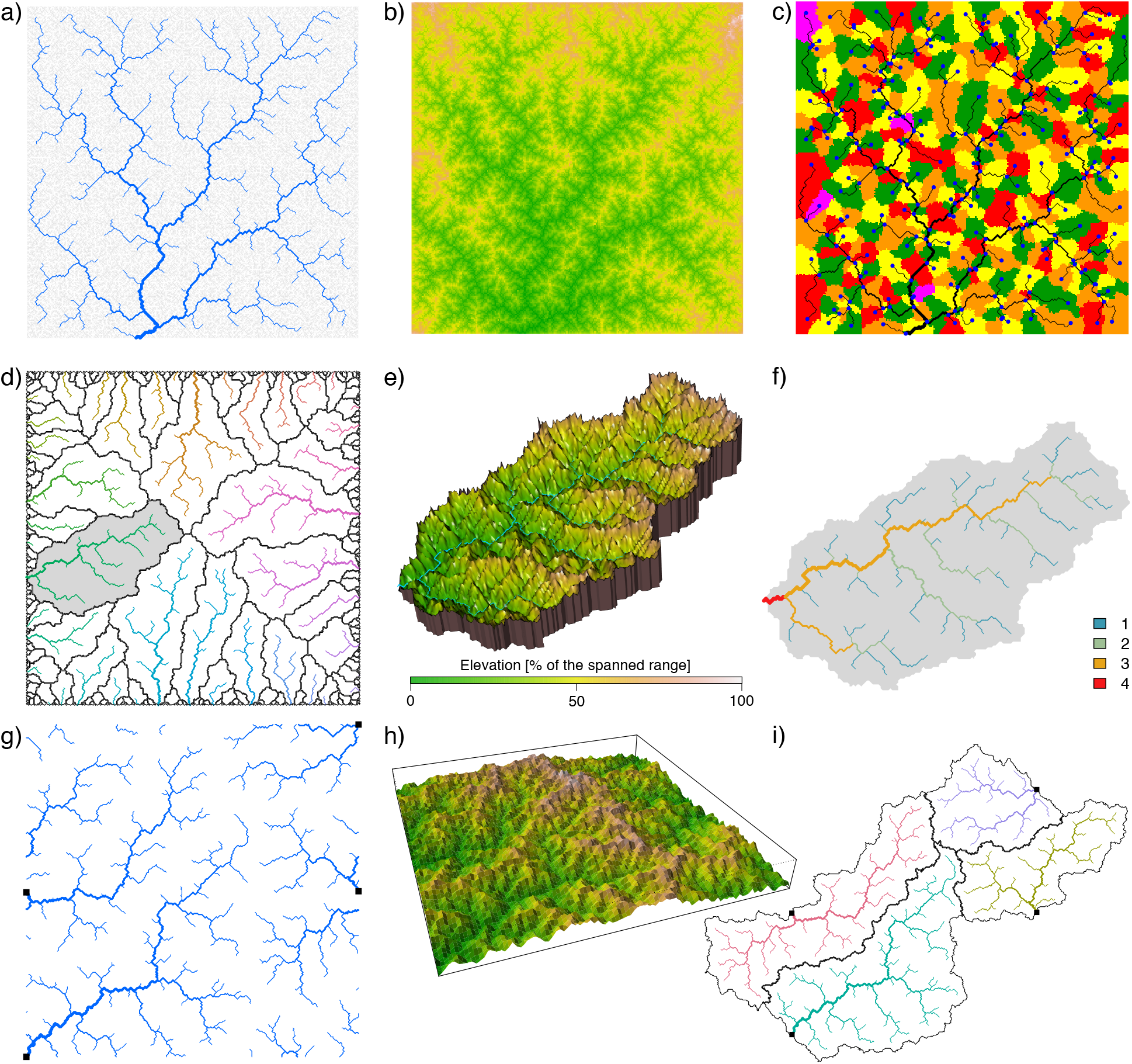
Examples of outputs from `OCNet’s` graphical functions. a) Representation of an OCN generated on a 250×250 lattice (`draw_simple_OCN`). Note that the network spans the whole lattice; for graphical reasons, the portion of network exceeding a given *A*_*T*_ is plotted in blue. b) Planar representation of the elevational landscape generated by the OCN of panel a (`draw_elev2D_OCN`). c) Partitioning of the lattice into subcatchments for the OCN of panel a (draw subcatchments OCN); blue dots indicate locations of the nodes at the AG level. d) Representation of an OCN generated on a 400×400 lattice, with all perimetral pixels as outlets (`draw_contour_OCN`); black solid lines display partitioning among catchments; the grey background identifies the largest catchment. e) 3D representation of the largest catchment within the OCN of panel d (draw elev3Drgl OCN). f) Strahler stream order values for the largest catchment within the OCN of panel d (`draw_thematic_OCN`). g) Representation of an OCN generated on a 300×300 lattice, with 4 outlets (shown by black squares) and periodic boundaries (`draw_contour_OCN`). h) Perspective 3D representation of the OCN of panel g (`draw_elev3D_OCN`). i) Real-shaped representation of the OCN of panel g (`draw_contour_OCN`). A script generating this figure can be found in the Supporting Information.

### 2.3 Aggregation levels

Before moving to the illustration of some possible applications of the package, we here clarify some concepts and terminology with respect to the aggregation of OCNs. Additional details are provided in the package documentation. Networks produced by the OCN algorithm can be used in a variety of fashions (see Table 1 for a review) by exploiting different connectivity metrics that are embedded in the OCN construct. At a first, non-aggregated level, each cell of the lattice (also termed as pixel) constitutes a node of the network (see Fig. 4), and the connectivity among nodes is ruled by the flow direction pattern (represented by the adjacency matrix) obeying the OCN principle. This is here referred to as the flow direction level (FD).

**Table 1:**
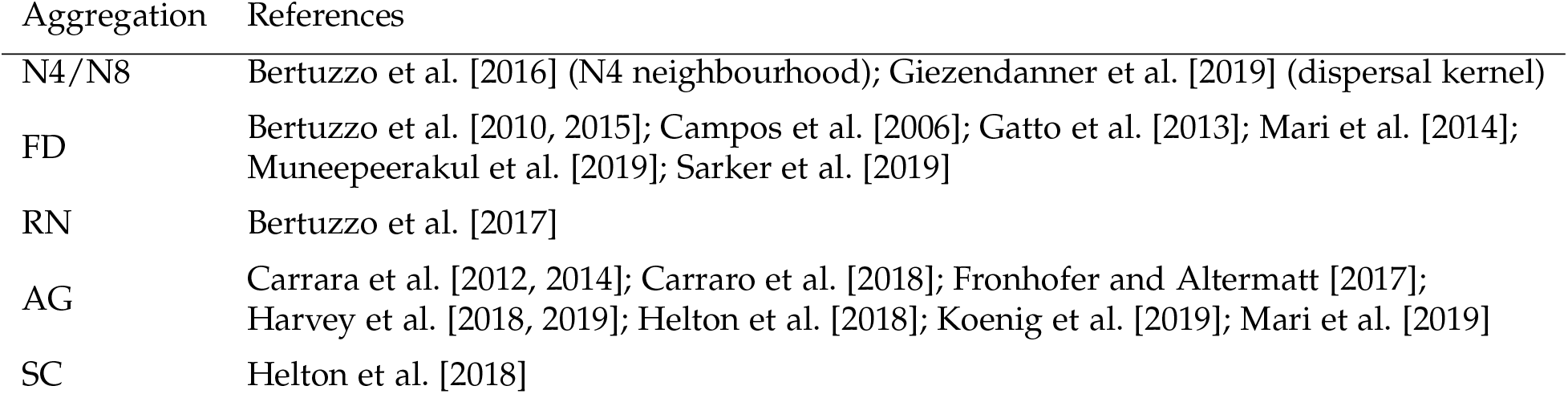
Types of OCN aggregation schemes used in previous studies.

**Figure 4:**
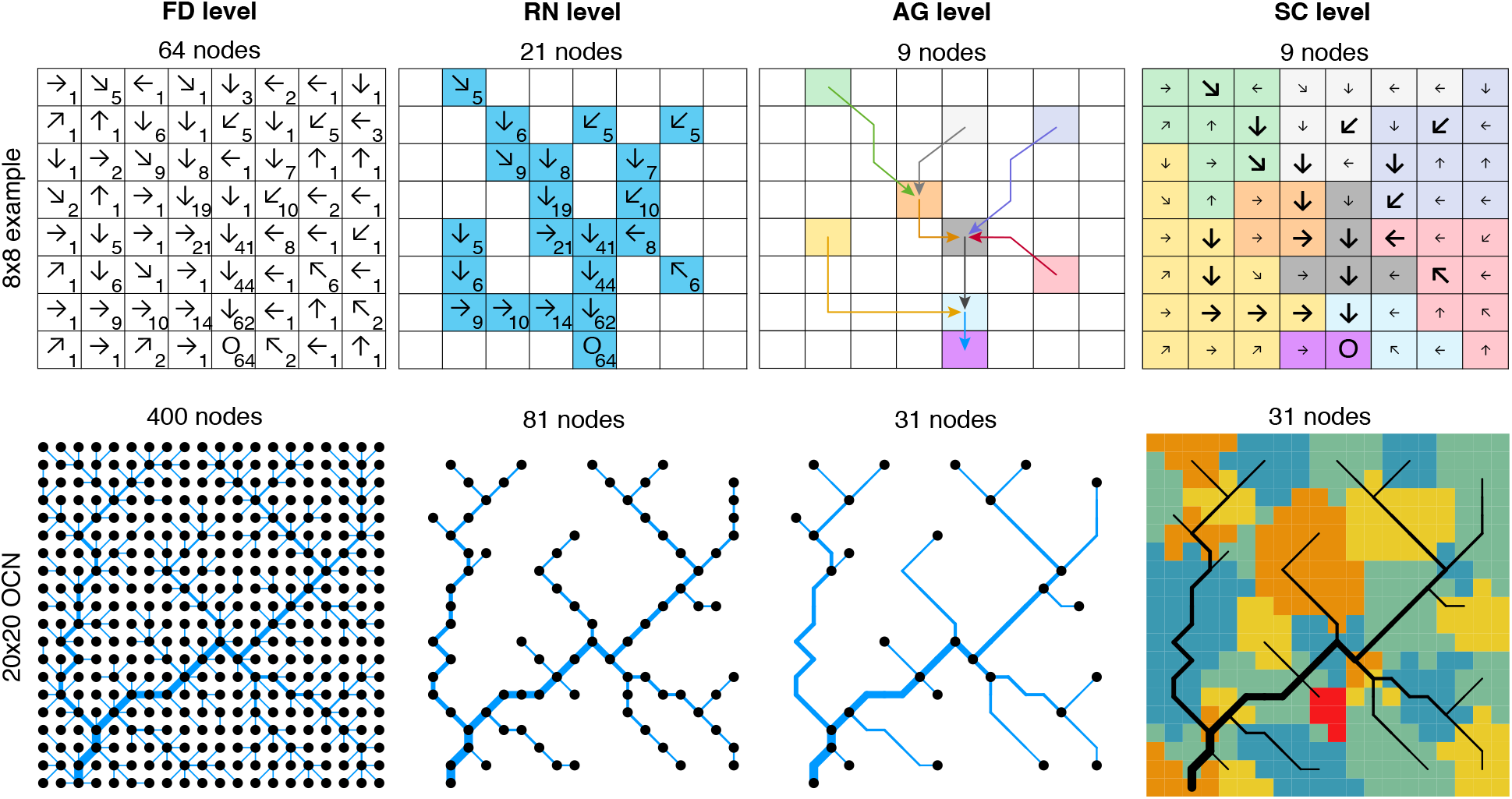
Representation of the different aggregation levels. Top row: example from a single-outlet 8×8 lattice describing how the aggregation procedure operated by `aggregate_OCN` works. Letter ‘O’ identifies the outlet pixel. Arrows on the other pixels identify flow directions; note that the these are not representative of an OCN, but are here presented only for illustrative purposes. Numbers represent the cumulative drainage area (in number of pixels). At the FD level, all 64 pixels belong to the network. To obtain the RN level, a threshold area *A*_*T*_ = 5 pixels is applied to distinguish pixels belonging to the river network. Bottom row: the same procedure is applied to a single-outlet, 20×20 OCN (obtained via the script presented in *Generation of an OCN*). Aggregation is performed with *A*_*T*_ = 5 pixels. Note that river width is proportional to the square root of drainage area [Leopold and Maddock, 1953]. A script generating this figure can be found in the Supporting Information.

As customary in hydrology when extracting a river network based on digital elevation models of the terrain [O’Callaghan and Mark, 1984], a threshold *A*_*T*_ on drainage area can be imposed to identify those pixels of the lattice that constitute nodes of the river network (RN, second level – see Terui et al. [2018] for an example of an ecological application of non-OCN synthetic networks akin to OCNs at the RN level). In a third, aggregated level (AG), nodes correspond to sources, confluences and outlet(s) of the river network identified at the RN level. The whole lattice is then partitioned into areas that directly drain into the nodes at the AG level, or the edges departing from them, thereby constituting the fourth, subcatchment level (SC).

A fifth level (catchment, CM) partitions the lattice into regions that are drained by different outlets, when the multiple-outlet option in create OCN is enabled (see following section). Finally, in an optional, zero-level spatial structure, all lattice pixels are treated as nodes, but connectivity follows the Von Neumann (4 nearest neighbours, level N4) or Moore (8 nearest neighbours, level N8) neighbourhoods, as in the green network described by Altermatt [2013]. In this case, the OCN was used to generate a realistic elevation gradient governed by fluvial erosion, on which, for instance, the structure of terrestrial metapopulations can be studied [Bertuzzo et al., 2016; Giezendanner et al., 2019].

The final output of the OCNet functions is a list of lists, each of which named after the corresponding aggregation level (N4/N8, FD, RN, AG, SC, CM) and containing relevant topological and morphological information for that level. Variables may vary in number, type and definition among the sub-lists, although the adjacency matrix is defined for all levels, while the drainage area vector is defined for all levels but N4/N8.

## 3 Overview of package features

### 3.1 Number of outlets, boundary types, and elevational gradients

Although the OCN principle is primarily intended to be applied to networks spanning the whole drainage domain (where the area drained by the outlet is equal to the lattice area, see an example in Fig. 3a–c), the generalization to the case of multiple networks—each of which subsumed by a different outlet—within the same lattice is straightforward. Indeed, the very same mathematical formulation presented in *Theoretical background* holds when multiple outlet pixels are imposed. Technically, this is done by preventing these pixels to drain into their neighbouring pixels. In this case, the sum of the areas drained by all outlet pixels is equal to the lattice area. In the package, the multiple-outlet option can be activated by means of the optional input nOutlet of function create OCN. In the limiting case, all pixels at the lattice boundary can be treated as outlets [Sun et al., 1994; Bertuzzo et al., 2017]. This is done by setting `nOutlet` = `“All”` in `create_OCN`. A graphical representation of an OCN obtained for the latter case is shown in Fig. 3d.

When a pixel’s flow direction is rewired during the search for an optimal network configuration, possible directions are generally those towards the eight neighbouring pixels. This is not the case for the outlet pixels (which cannot be rewired) and the pixels at the lattice boundaries, which can be rewired to either three (corner pixels) or five (side pixels) neighbouring cells. This latter assumption can be relaxed by allowing pixels at the boundary to drain into eight neighbours, by also considering pixels at the opposite sides as feasible directions. In `OCNet`, periodic boundaries can be enabled via the optional input `periodicBoundaries of create OCN`. An example is shown in Fig. 3g–i. Such option can be useful when the OCN lattice is to be considered as the periodic unit of an infinite landscape [Bertuzzo et al., 2016; Giezendanner et al., 2019], or when one aims at building OCNs spanning domains that are not lattice-shaped (see Fig. 3f, i).

Once an OCN has been created by the simulated annealing algorithm, the iterative application of the slope-area relationship starting from the outlet node and moving in the upstream directions enables the derivation of the elevation field subsumed by the OCN (up to two constants, e.g., the elevation and slope of the outlet pixel). Some examples of elevational landscapes built on OCNs are provided in Fig. 3b, e, h. Importantly, the slope-area relationship only holds for the channeled portion of the domain, which implies, strictly speaking, that the OCN must not be aggregated if one aims at making use of a three-dimensional landscape generated by an OCN. Moreover, the slope-area relationship is actually multiscaling [Tarboton et al., 1989], therefore the simple recursive application of 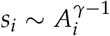 (as performed by the function `landscape_OCN`) to yield an elevational landscape is to be considered as a first approximation, suitable for ecological applications. Methods to account for the scaling of the variance of the slope-area relationship exist [Grimaldi et al., 2005], but are beyond the scope of this work.

### 3.2 Relationship between threshold drainage area and number of nodes

Owing to the somewhat heuristic procedure for the definition of an aggregated network based on a threshold drainage area value *A*_*T*_, it is not possible to establish a priori how many nodes at the AG level correspond to a given *A*_*T*_. This in fact depends on the configuration of the OCN at the FD level, which is the result of a stochastic process. This issue is particularly relevant when OCNs are used in experiments where practical reasons enforce a limitation on the number of nodes that can be handled, or when several OCN replicas with the same number of aggregated nodes are required.

To help overcome this issue, OCNet includes the function `find area threshold OCN`, which requires as input a non-aggregated OCN (produced by `landscape_OCN`) and evaluates the number of nodes resulting from the aggregation procedure for different values of *A*_*T*_. Such a function can therefore be used prior to `aggregate_OCN` to assess which threshold has to be used to obtain a network with the desired aggregation structure. Additionally, `find_ area_threshold_OCN` also evaluates other variables that help characterize the network structure from a hydrological perspective, such as maximum stream order and drainage density. Maximum stream order can be of interest in some studies, when patch sizes need to be related to the structure of the underlying network but only few discrete dimensions are available, so that it is convenient to employ different patch sizes for different stream order values of the corresponding nodes (e.g., Harvey et al. [2018]). Drainage density is relevant because it allows the assessment of hydrological characteristics of the aggregated OCN for a given metric resolution (i.e., the length in meters attributed to a pixel length—corresponding to the optional input `cellsize of create_OCN`), such as aridity and timing of the hydrologic response [Pallard et al., 2009].

Fig. 5 shows results from the application of `find_area_threshold_OCN` to several OCNs built on large lattices. When *A*_*T*_ is lower than 2% of the lattice size, the number of nodes scales fairly well as a power law of the normalized threshold area (Fig. 5a). This relationship allows qualitatively assessing the relevant range of *A*_*T*_ corresponding to a sought number of nodes at the AG level, which can be used as input in `find_area_threshold_OCN` to speed up its execution, especially for large networks. Scaling relationships with *A*_*T*_ are also found for maximum Strahler stream order (Fig. 5b) and drainage density (Fig. 5c). To provide an example, if a threshold *A*_*T*_ = 20 pixels is applied to a 200×200 OCN, the expected number of nodes at the AG level is 1052.4, the expected maximum stream order is 5.56, and the expected drainage density is *D*_*d*_ = 0.1454 inverse planar units, which corresponds to a relatively wet catchment (*D*_*d*_ = 2.91 km^−1^) of area 100 km^2^ (when 1 planar unit represents 50 m), or to a rather arid catchment (*D*_*d*_ = 1.46 km^-1^) of area 400 km^2^ (if 1 planar unit is equal to 100 m). Notably, the relationship between drainage density and threshold area *A*_*T*_ mirrors the scaling behaviour of drainage areas (Fig. 5d), which is characterized by an exponent *β* in the range [0.42; 0.45] (see Introduction and Rinaldo et al. [2014]). Indeed, drainage density for a given *A*_*T*_ is roughly (i.e., if differences in lengths between vertical/horizontal and diagonal flow directions are neglected) equal to the number of pixels whose area is greater than or equal to *A*_*T*_. Fig. 5d also represents how the number of nodes at the RN level (*N*_*RN*_) scales with varying *A*_*T*_: to this end, it suffices to replace *a* with *A*_*T*_ and *P*(*A* ≥ *a*) with *N*_*RN*_ /*N*.

**Figure 5:**
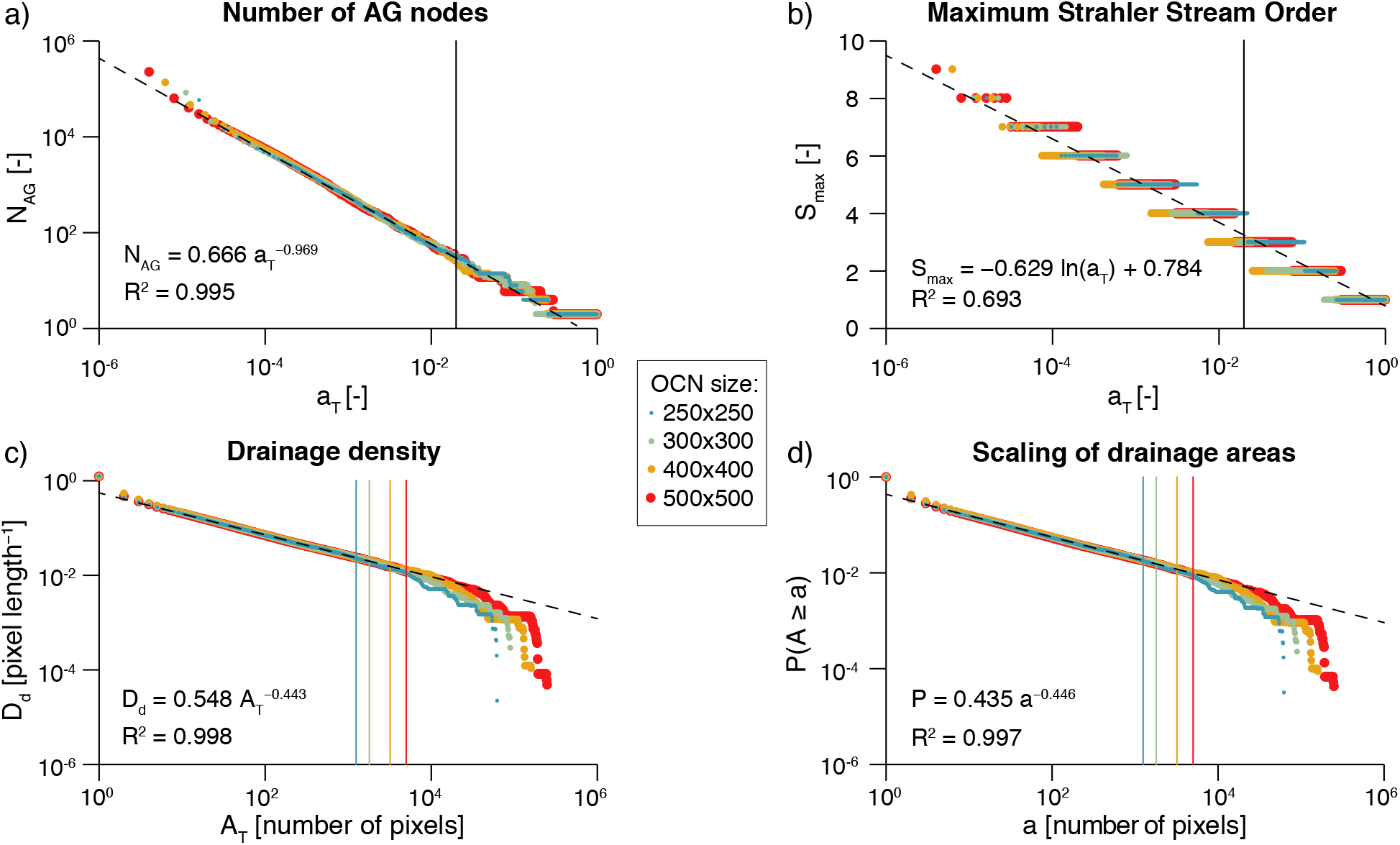
Effect of choice of threshold area *AT* on OCN configuration. Aggregation of four large OCNs is performed ∀*AT* = {1, …, *N*} via function `find_area_threshold_OCN`. For all panels, vertical lines indicate the cutoff value *A_T_* = 0.02 *N*; only points corresponding to threshold area values below the cutoff are used to estimate the (dashed) regression lines, whose equations and R^2^ values are reported. a) Number of nodes at the AG level scales as a power-law function of the normalized threshold area a_T_ = *A*_*T*_ /*N*. b) Maximum Stahler stream order value as a function of normalized threshold area a_T_. c) Drainage density scales as a power-law function of threshold area *A*_*T*_. d) Scaling behaviour of OCNs: probability *P*(*A ≥ a*) of randomly sampling a pixel within the lattice whose drainage area *A* is not greater than a given value a. A script generating this figure can be found in the Supporting Information.

The scaling behaviour of OCNs displayed in Fig. 5 can also provide useful information with respect to the choice of values of relevant parameters *N* and *A*_*T*_ that allow generating an OCN of adequate size for practical applications. To this extent, it is worthwhile to note that the OCN construct is invariant under coarse graining [Rodriguez-Iturbe and Rinaldo, 2001; Rinaldo et al., 2014], which means that the choice of the lattice dimension *N* does not affect the scaling of drainage areas. As in the example above, such choice should rather be based on geomorphological arguments, that is, the answers to the questions: What is the area that the OCN is supposed to drain? What are the expected values of maximum stream order and drainage density on this area? However, as a rule-of-thumb indication, we suggest to perform aggregation with a threshold not greater than *A*_*T*_ = 0.02 · *N*, such that the obtained configuration is not affected by the finite-size scaling effect; this corresponds to an expected *N*_*AG*_ ≥ 30 (see Fig. 5a).

### 3.3 Compatibility with existing R-packages

Specific functions of `OCNet` enable transformation of OCNs into objects that can be used by other commonly used R-packages in spatial ecology. In particular, compatibility with igraph [Csardi and Nepusz, 2006], a package for network analysis and visualization, is provided by function `OCN_to_igraph`. More-over, function `OCN_to_SSN` transforms an OCN at a desired aggregation level into an object that can be read by `SSN`, a package on spatial statistical modeling and prediction for data on stream networks [Ver Hoef et al., 2014]. Examples for these functions are shown in Fig. 6. Finally, output from OCNet can be used in combination with R-packages for geostatistical modelling such as gstat [Pebesma, 2004], based on the coordinates of nodes of an OCN given at any aggregation level. Remarkably, adjacency matrices and other information can easily be extracted as base R objects, which guarantees compatibility with virtually every R-package and even other programming languages.

**Figure 6:**
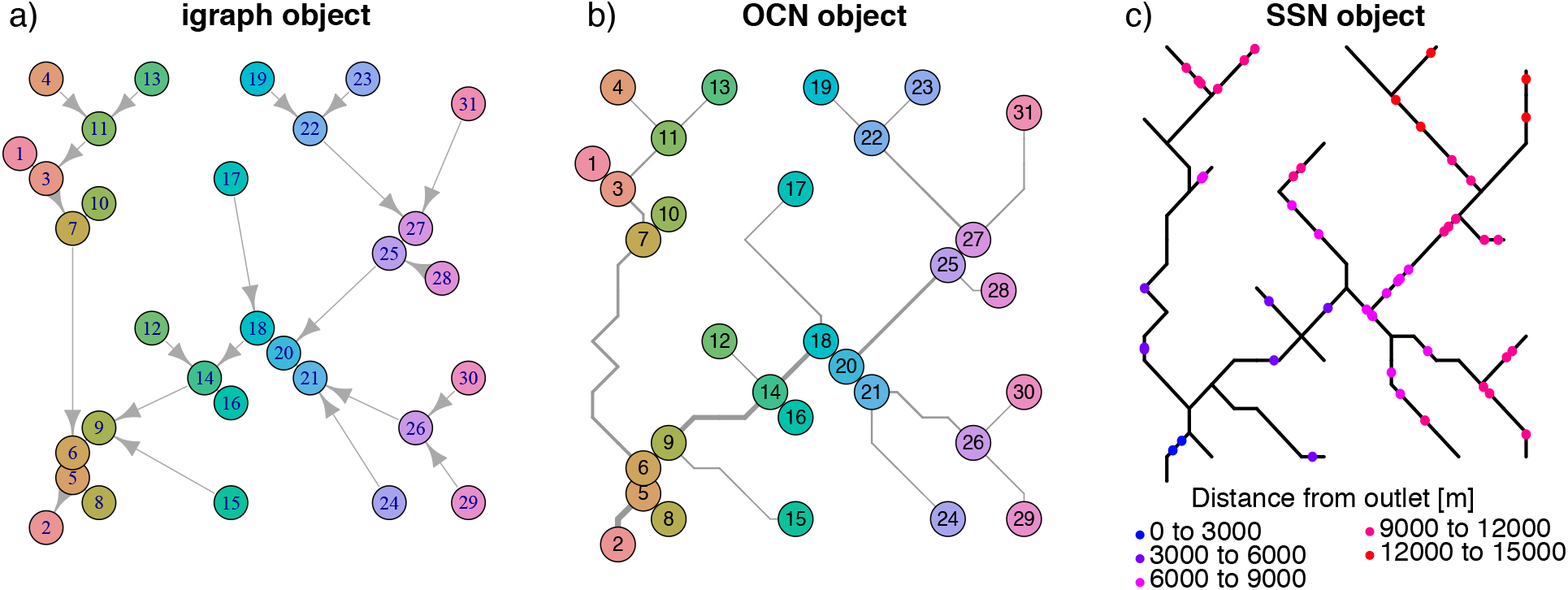
Compatibility of `OCNet` with packages igraph and `SSN`. Examples are built on the OCN obtained in *Generation of an OCN*. a) The OCN, aggregated at the AG level, is transformed into an igraph object (via `OCN_to_igraph`), and plotted via the function `plot.igraph of igraph`. b) The same OCN is plotted via `draw_thematic_OCN`. c) The same OCN is transformed into an SSN object (via `OCN_to_SSN`) and plotted via the function `plot.SpatialStreamNetwork of SSN`. Here, 50 observation points have been sampled along the network by means of a binomial design, and their distance from the outlet is displayed. A script generating this figure can be found in the Supporting Information.

## 4 Application example: a metapopulation model

In order to show a possible application of the `OCNet` package, we now apply a simple metapopulation model for an invasive riverine species to an OCN.

### 4.1 Generation of an OCN

Let us build an OCN with the following assumptions: it spans a 20×20 lattice, with a single outlet located close to the southwestern corner of the lattice, and each pixel represents a square of side 500 m (total size of the catchment is therefore 100 km^2^); the elevation, slope and channel width of the outlet node are 0 m a.s.l., 0.01, and 5 m, respectively; the threshold area used to aggregate the network is equal to 1.25 km^2^. The code lines to build such network are the following:

~~~
set.seed(1) # use fixed random number generator
OCN <- create_OCN(20, 20, outletPos = 3, cellsize = 500)
OCN <- landscape_OCN(OCN, slope0 = 0.01)
OCN <- aggregate_OCN(OCN, thrA = 1.25e6)
OCN <- paths_OCN(OCN, pathsRN = TRUE)
OCN <- rivergeometry_OCN(OCN, widthMax = 5)
~~~

The resulting OCN is shown in Fig. 4.

### 4.2 Metapopulation model

Let us build a discrete-time, deterministic metapopulation model on the previously built OCN, according to the following assumptions: (i) the model is run on the OCN aggregated at the RN level (consisting of *N*_*n*_ nodes); (ii) population growth at each node follows the Beverton-Holt model [Beverton and Holt, 1957], with baseline fecundity rate *r* = 1.05 constant for all nodes, and carrying capacity *K*_*i*_ = 10 · *W*_*i*_, where *W*_*i*_ is the river width of the network node *i*; (iii) at each time step *t*, the number of individuals moving from node *i* is equal to *gP*_*i*_(*t*), where *g* = 0.1 is a mobility rate constant for all nodes, and *P*_*i*_(*t*) is the (expected) population size at node *i* and time *t*; (iv) at each time step, individuals at node *i* can only move to a node that is directly connected to *i*, either downstream or upstream; (v) *p*_*d*_ and *p*_*u*_ = 1 − *p*_*d*_ identify the probability to move downstream or upstream, respectively; (vi) if the indegree of node *i* is larger than 1 (namely the node has multiple upstream connections), individuals moving upstream are split among the possible destination nodes into fractions *ϒ*_*i*_ proportional to their drainage areas; (vii) as initial condition, all network nodes are uninhabited barring the node *f* that is farthest from the outlet (where *P*_*f*,1_ = 1). The model equation hence reads

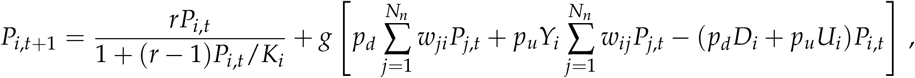

where *w*_*ij*_ is a generic entry of the adjacency matrix **W** expressing OCN connectivity at the RN level; *D*_*i*_ (*U*_*i*_) is equal to one if there is a downstream (upstream) connection available from node *i* and is null otherwise. Weights *ϒ*_*i*_ are defined as

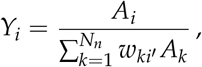

where *i*′ is such that *w_ii_*′ = 1, and *A*_*i*_ is the drainage area at node *i*. A code script performing model simulations is reported in the Supporting Information.

We performed two model simulations to investigate the effect of parameters *p*_*d*_, *p*_*u*_ on the time elapsed until the system reaches a steady state; in a first (default) run, no preferential direction of movement was assumed (*p*_d_ = *p*_*u*_ = 0.5); in the second run, a preference for downstream movement (*p*_*d*_ = 0.7, *p*_*u*_ = 0.3) was hypothesized. Results are shown in Fig. 7. When *p*_*d*_ = 0.5, the invading species rapidly reaches the equilibrium in the initially occupied node, while colonization of the downstream patches is delayed. When a preference for downstream movement is attributed (*p*_*d*_ = 0.7), local population growth in the onset (green) node is slower, whereas invasion of the outlet node occurs faster, both in terms of initial growth and establishment of the equilibrium (see coloured vertical lines in Fig. 7a). Colonization of the headwater that is farthest from the onset node is also delayed with respect to the default case. As a result, when *p*_*d*_ = 0.7, the metapopulation size initially grows faster than when *p*_*d*_ = 0.5, due to fast invasion of the downstream nodes and growth of the local populations therein (see Fig. 7b); in a second phase, the growth rate of the metapopulation is reduced, because invasion of the upstream nodes is hampered by the low *p*_*u*_ value, and the establishment of the equilibrium is delayed. As for the spatial spread of the metapopulation, when a preference for downstream movement is adopted, local population sizes at equilibrium tend to increase in the downstream nodes and decrease in the upstream nodes with respect to the default case (Fig. 7a), resulting in a slightly lower overall metapopulation size at equilibrium (Fig. 7b).

**Figure 7:**
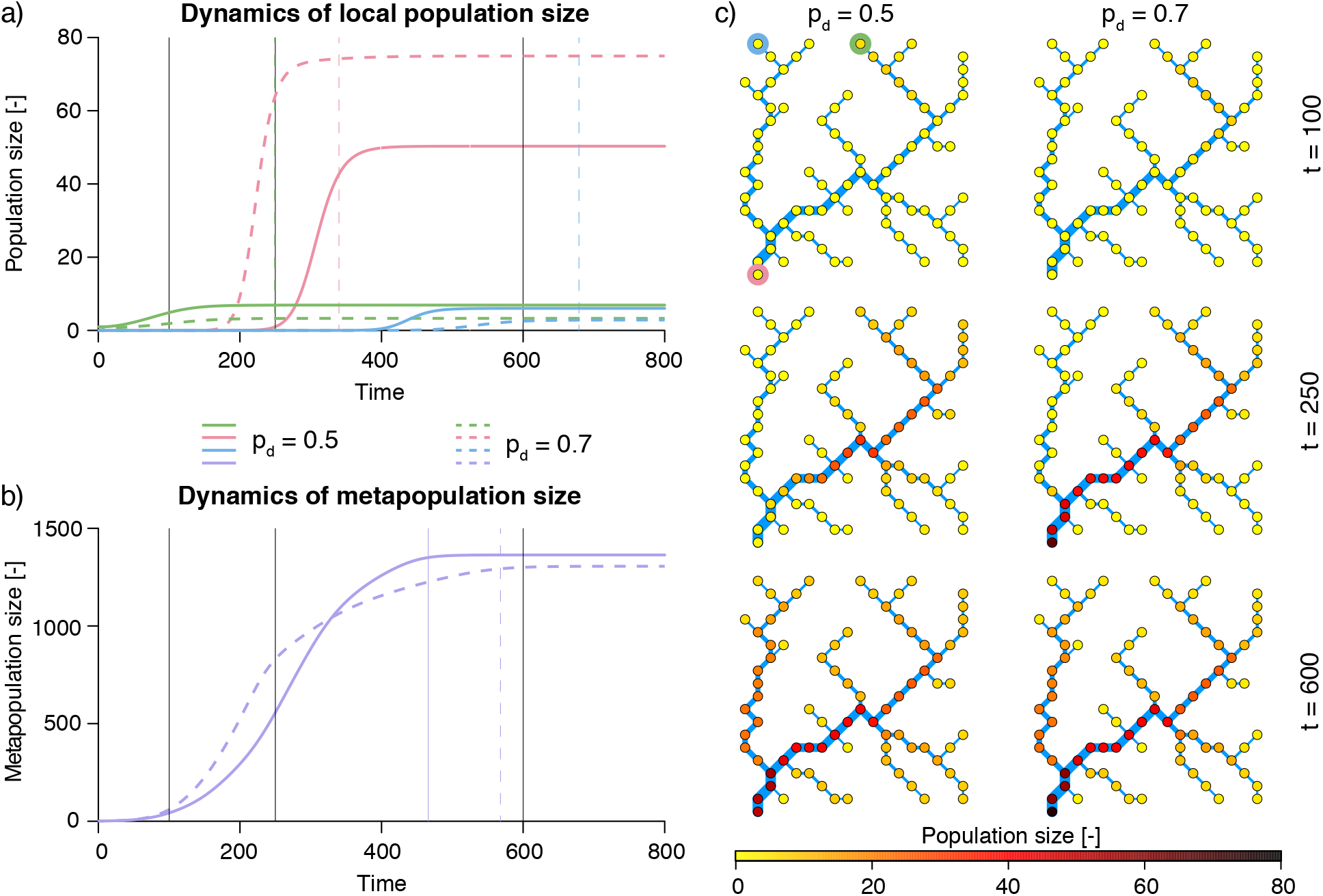
Results from the application of a metapopulaton model on an OCN. a) Dynamics of local population size in three nodes: green represents the headwater that is invaded at the beginning of the simulation; red, the outlet; blue, the headwater that is farthest from the green node (see circles in the top-left corner of panel c). Coloured vertical lines represent the time steps when the respective local population has reached the equilibrium (arbitrarily imposed as 99% of the population size at *t* = 800). Black solid lines identify the time steps used in panel c. b) Dynamics of the overall metapopulation size. Line styles as in panel a. c) Snapshots (obtained via `draw_thematic_OCN`) of spatial spread of the metapopulation at 3 different time steps. A script generating this figure can be found in the Supporting Information.

## 5 Conclusions

The importance of adequately representing spatial processes in ecological and evolutionary studies cannot be overstated. In the realm of freshwater ecology in particular, it is essential to consider how geomorphology shapes the structure of dendritic river networks and the ensuing connectivity configuration, which in turn control the variability of physical habitats and environmental variables, the dispersal of species and pathogens, and the spatial patterns of biodiversity and ecosystem processes. To this end, we presented `OCNet`, an R-package that enables the generation of optimal channel networks, spanning trees that reproduce all scaling features of real river networks throughout the globe. These can be used as realistic riverine landscape analogues for a number of ecological, epidemiological, ecohydrological and evolutionary studies. We reviewed the theoretical background of the OCN concept and the existing applications on problems of ecological relevance, provided an overview of the main functionalities of the package, and proposed an example of application in the context of an invasive riverine species. We believe that this tool will allow a leap forward in the way spatial processes in river networks are investigated.

## Acknowledgements

FA acknowledges funding from the Swiss National Science Foundation Grants No PP00P3 179089 and 31003A 173074 and the University of Zurich Research Priority Programme “URPP Global Change and Biodiversity”. This is publication ISEM-YYYY-XXX of the Institut des Sciences de l’Evolution – Montpellier.

## Authors contributions

LC wrote the first draft of the manuscript, and developed the package based on preliminary scripts from EB with inputs and contributions from EAF, RF, and IG. EB, AR, and FA conceived the study. All authors contributed to the final version of the manuscript.

## Data accessibility

The code of the `OCNet` package is accessible on both CRAN (https://CRAN.R-project.org/package=OCNet) and Github (http://doi.org/10.5281/zenodo.3669873 - development version).

