## Supporting code and data for "Generation and application of river network analogues for use in ecology and evolution": figures_OCNet_manuscript.R

```
# Code reproducing figures of manuscript
# "Generation and application of river network analogues for use in ecology and evolution"
# by Carraro Luca et al.
# uploaded on bioRxiv

#### PACKAGE INSTALLATION AND LIBRARY LOADING ####
rm(list=ls())
setwd("C:/Users/carrarlu/Dropbox/Applicazioni/ShareLaTeX/OCNet-MEE/code")

# install devtools (for following installations)
if (!require(devtools)){
  install.packages("devtools")}

# install OCNet
if (!require(OCNet)){
  devtools::install_github("lucarraro/OCNet")}

# install igraph
if (!require(igraph)){
  install.packages("igraph")}

# install SSN
if (!require(SSN)){
  install.packages("SSN")}

# install extended OCNet dataset from Github (needed for Fig. 2)
# the extended dataset contains "energy" vectors for the ready-made OCNs
# this might take some minutes
if (!require(OCNetExtendedData)){
  devtools::install_github("lucarraro/OCNetExtendedData")}

library(devtools)
library(OCNet)
library(OCNetExtendedData)
library(igraph)
library(SSN)

old.par <- par()

# # # # # # # # # # # # # # # # # # # # # # # # # # # # # # # # # 
#### FIGURE 2 - EFFECT OF INITIAL STATE AND COOLING SCHEDULE ####
# # # # # # # # # # # # # # # # # # # # # # # # # # # # # # # # # 

cat("Creating Figure 2...\n")

OCN0 <- create_OCN(25,25,nIter=0)
OCN_V0 <- create_OCN(25,25,typeInitialState="V",nIter=0)

colors <- hcl.colors(13,palette="Zissou 1")

subset_iter <- seq(100,(length(OCN_250$energy)-1),100)

nIter <- OCN_250$nIter
nNodes <- OCN_250$dimX*OCN_250$dimY

TemperatureHot <- c(1+numeric(OCN_250_hot$initialNoCoolingPhase*nIter),
                    exp(-OCN_250_hot$coolingRate*(1:(nIter-OCN_250_hot$initialNoCoolingPhase*nIter))/nNodes))
TemperatureCold <- c(1+numeric(OCN_250_cold$initialNoCoolingPhase*nIter),
                    exp(-OCN_250_cold$coolingRate*(1:(nIter-OCN_250_cold$initialNoCoolingPhase*nIter))/nNodes))
TemperatureDef <- c(1+numeric(OCN_250$initialNoCoolingPhase*nIter),
                    exp(-OCN_250$coolingRate*(1:(nIter-OCN_250$initialNoCoolingPhase*nIter))/nNodes))

## Plot figure
par(mfrow=c(1,1),mai=c(1,1,1,1))
plot(subset_iter,OCN_250_hot$energy[subset_iter]/OCN_250_hot$energy[1], type="l", col=colors[13], 
     ylim=c(0,1),bty="n",xaxt="n",yaxt="n",xlab="iterations",ylab="Energy, Temperature")
axis(1,pos=0); axis(2,pos=0)
lines(subset_iter,OCN_250_V_hot$energy[subset_iter]/OCN_250_V_hot$energy[1],col=colors[11])
lines(subset_iter,OCN_250$energy[subset_iter]/OCN_250$energy[1],col=colors[8])
lines(subset_iter,OCN_250_V$energy[subset_iter]/OCN_250_V$energy[1],col=colors[6])
lines(subset_iter,OCN_250_cold$energy[subset_iter]/OCN_250_cold$energy[1], col=colors[3])
lines(subset_iter,OCN_250_V_cold$energy[subset_iter]/OCN_250_V_cold$energy[1], col=colors[1])
lines(subset_iter,TemperatureHot[subset_iter],lty=2,col=colors[12])
lines(subset_iter,TemperatureDef[subset_iter],lty=2,col=colors[7])
lines(subset_iter,TemperatureCold[subset_iter],lty=2,col=colors[2])

par(mfrow=c(2,4), mai=c(0,0,0,0))
draw_simple_OCN(OCN_250_cold,riverColor = colors[3])
draw_simple_OCN(OCN_250,riverColor = colors[8])
draw_simple_OCN(OCN_250_hot,riverColor = colors[13])
draw_simple_OCN(OCN0,riverColor = "#909090",thrADraw = 0)
draw_simple_OCN(OCN_250_V_cold,riverColor = colors[1])
draw_simple_OCN(OCN_250_V,riverColor = colors[6])
draw_simple_OCN(OCN_250_V_hot,riverColor = colors[11])
draw_simple_OCN(OCN_V0,riverColor = "#909090",thrADraw = 0)


par(old.par)

# # # # # # # # # # # # # # # # # # # # # # # #
#### FIGURE 3 - TESTING PLOTTING FUNCTIONS ####
# # # # # # # # # # # # # # # # # # # # # # # #

cat("Creating Figure 3...\n")

## Load data
if (!(file.exists("data_fig3.rda"))){
  # NB: Running this chunk of code takes some minutes
  OCN <- OCN_250_V_hot
  OCN <- landscape_OCN(OCN)
  OCN <- aggregate_OCN(OCN)
  
  OCN2 <- OCN_300_4out_PB_hot
  OCN2 <- landscape_OCN(OCN2, optimizeDZ = T, displayUpdates = 2)
  
  OCN3 <- OCN_400_Allout
  OCN3 <- landscape_OCN(OCN3, displayUpdates = 2, slope0=0.01)
  OCN3 <- aggregate_OCN(OCN3, thrA=2e5)
  
  save("OCN","OCN2","OCN3",file="data_fig3.rda")
  
} else {load("data_fig3.rda")}

## Plot figure
par(mfrow=c(3,3), mai=c(0.1,0.1,0.1,0.1))

draw_simple_OCN(OCN, easyDraw = FALSE)
draw_elev2D_OCN(OCN)
draw_subcatchments_OCN(OCN)
points(OCN$AG$X,OCN$AG$Y,pch=19,col="blue",asp=1, cex=0.25)

draw_contour_OCN(OCN3)
plot(0,0,col=NULL,axes=FALSE)
draw_thematic_OCN(OCN3$AG$streamOrder,OCN3,colPalette=hcl.colors(4, palette="Zissou 1"),
                  discreteLevels=TRUE,chooseCM=TRUE)
draw_elev3Drgl_OCN(OCN3,chooseCM=TRUE, aspect=c(1,1,0.1), drawRiver = TRUE)

draw_contour_OCN(OCN2, drawContours = FALSE, drawOutlets = 2, exactDraw = FALSE, colPalRiver = "#0066ff")
draw_elev3D_OCN(OCN2, drawRiver = F, addColorbar = F, coarseGrain = c(3,3), expand=0.1)
draw_contour_OCN(OCN2, drawOutlets = 2)

par(old.par)

# # # # # # # # # # # # # # # # # # # # # # # #
#### FIGURE 4 - AGGREGATION LEVELS EXAMPLE ####
# # # # # # # # # # # # # # # # # # # # # # # #

cat("Creating Figure 4...\n")

## Build OCN
set.seed(1)
OCN <- create_OCN(20, 20, outletPos = 3, cellsize = 500)
OCN <- landscape_OCN(OCN, slope0 = 0.01)

OCN <- aggregate_OCN(OCN, thrA = 5*500^2)
OCN0 <- aggregate_OCN(OCN, thrA = 0) # non-aggregated OCN
# NB: Reply RN or AG to the question appearing in the console

## Plot Figure
par(mfrow=c(1,4))
draw_thematic_OCN(numeric(OCN0$FD$nNodes),OCN0,drawNodes=TRUE, chooseAggregation = "RN",
                  backgroundColor = NULL,cex = 1,discreteLevels = T,
                  colPalette="black",addLegend = F)

title(sprintf("%d nodes",OCN0$RN$nNodes))
draw_thematic_OCN(numeric(OCN$RN$nNodes),OCN,drawNodes=TRUE,
                  backgroundColor = NULL,cex = 1,discreteLevels = T,
                  colPalette="black",addLegend = F)
title(sprintf("%d nodes",OCN$RN$nNodes))
draw_thematic_OCN(numeric(OCN$AG$nNodes),OCN,drawNodes=TRUE,
                  backgroundColor = NULL,cex = 1,discreteLevels = T,
                  colPalette="black",addLegend = F)
title(sprintf("%d nodes",OCN$AG$nNodes))
draw_subcatchments_OCN(OCN,colPalette=hcl.colors(5,palette="Zissou 1"))
title(sprintf("%d nodes",OCN$SC$nNodes))

par(old.par)

# # # # # # # # # # # # # # # # # # # # # # # #
#### FIGURE 5 - THRESHOLD AREA AND SCALING ####
# # # # # # # # # # # # # # # # # # # # # # # #

cat("Creating Figure 5...\n")

### Load data 
if (!(file.exists("data_fig5.rda"))){
  # NB: Running this chunk of code takes some hours
  OCN <- landscape_OCN(OCN_250,displayUpdates = 2)
  thr250 <- find_area_threshold_OCN(OCN,displayUpdates = 1)
  a250 <- numeric(OCN$FD$nNodes)
  P250 <- numeric(OCN$FD$nNodes)
  for (i in 1:OCN$FD$nNodes){
    a250[i] <- i*OCN$cellsize^2
    P250[i] <- sum(OCN$FD$A>=a250[i])/OCN$FD$nNodes
  } 
  
  OCN <- landscape_OCN(OCN_300_diag,displayUpdates = 2)
  thr300 <- find_area_threshold_OCN(OCN,displayUpdates = 1)
  a300 <- numeric(OCN$FD$nNodes)
  P300 <- numeric(OCN$FD$nNodes)
  for (i in 1:OCN$FD$nNodes){
    a300[i] <- i*OCN$cellsize^2
    P300[i] <- sum(OCN$FD$A>=a300[i])/OCN$FD$nNodes
  } 
  
  OCN <- landscape_OCN(OCN_400_T_hot,displayUpdates = 2)
  thr400 <- find_area_threshold_OCN(OCN,displayUpdates = 1)
  OCN <- OCN_400_T_hot
  a400 <- numeric(OCN$FD$nNodes)
  P400 <- numeric(OCN$FD$nNodes)
  for (i in 1:OCN$FD$nNodes){
    a400[i] <- i*OCN$cellsize^2
    P400[i] <- sum(OCN$FD$A>=a400[i])/OCN$FD$nNodes
  } 
  
  OCN <- landscape_OCN(OCN_500_hot,displayUpdates = 2)
  thr500 <- find_area_threshold_OCN(OCN,displayUpdates = 1)
  OCN <- OCN_500_hot
  a500 <- numeric(OCN$FD$nNodes)
  P500 <- numeric(OCN$FD$nNodes)
  for (i in 1:OCN$FD$nNodes){
    a500[i] <- i*OCN$cellsize^2
    P500[i] <- sum(OCN$FD$A>=a500[i])/OCN$FD$nNodes
  } 
  
  save(thr250,thr300,thr400,thr500,a250,P250,a300,P300,a400,P400,a500,P500,file="data_fig5.rda")
  
} else {load("data_fig5.rda")}

### Perform binning
# Figure would be too heavy if all points were displayed. Therefore, all vectors are binned into 1000 components.
# Min spacing is logarithmic. For each bin, mean number of nodes and mean drainage density are displayed.
# Mode function is used to compute maximum stream order for every bin (non-integer values would arise with mean)
# Note that regression lines refer to the whole vectors, not to the binned ones

nbins <- 1000

bin_thr250 <- seq(log10(min(thr250$thrValues)/max(thr250$thrValues)),0,length.out = nbins + 1)
thr_250_b <- nN_250_b <- SO_250_b <- DD_250_b <- numeric(nbins)

## define mode function
Mode <- function(x) {
  ux <- unique(x)
  ux[which.max(tabulate(match(x, ux)))]
}

for (i in 1:nbins){
  if (i < nbins){
    tmp <- which(log10(thr250$thrValues/max(thr250$thrValues)) >= bin_thr250[i] & 
                   log10(thr250$thrValues/max(thr250$thrValues)) < bin_thr250[i+1])
  } else {
    tmp <- which(log10(thr250$thrValues/max(thr250$thrValues)) >= bin_thr250[i] & 
                   log10(thr250$thrValues/max(thr250$thrValues)) <= bin_thr250[i+1])  
  }
  thr_250_b[i] <- mean(thr250$thrValues[tmp])
  nN_250_b[i] <- mean(thr250$nNodesAG[tmp])
  SO_250_b[i] <- Mode(thr250$streamOrder[tmp])
  DD_250_b[i] <- mean(thr250$drainageDensity[tmp])
}

bin_thr300 <- seq(log10(min(thr300$thrValues)/max(thr300$thrValues)),0,length.out = nbins + 1)
thr_300_b <- nN_300_b <- SO_300_b <- DD_300_b <- numeric(nbins)
for (i in 1:nbins){
  if (i < nbins){
    tmp <- which(log10(thr300$thrValues/max(thr300$thrValues)) >= bin_thr300[i] & 
                   log10(thr300$thrValues/max(thr300$thrValues)) < bin_thr300[i+1])
  } else {
    tmp <- which(log10(thr300$thrValues/max(thr300$thrValues)) >= bin_thr300[i] & 
                   log10(thr300$thrValues/max(thr300$thrValues)) <= bin_thr300[i+1]) 
  }
  thr_300_b[i] <- mean(thr300$thrValues[tmp])
  nN_300_b[i] <- mean(thr300$nNodesAG[tmp])
  SO_300_b[i] <- Mode(thr300$streamOrder[tmp])
  DD_300_b[i] <- mean(thr300$drainageDensity[tmp])
}

bin_thr400 <- seq(log10(min(thr400$thrValues)/max(thr400$thrValues)),0,length.out = nbins + 1)
thr_400_b <- nN_400_b <- SO_400_b <- DD_400_b <- numeric(nbins)
for (i in 1:nbins){
  if (i < nbins){
    tmp <- which(log10(thr400$thrValues/max(thr400$thrValues)) >= bin_thr400[i] & 
                   log10(thr400$thrValues/max(thr400$thrValues)) < bin_thr400[i+1])
  } else {
    tmp <- which(log10(thr400$thrValues/max(thr400$thrValues)) >= bin_thr400[i] & 
                   log10(thr400$thrValues/max(thr400$thrValues)) <= bin_thr400[i+1])  
  }
  thr_400_b[i] <- mean(thr400$thrValues[tmp])
  nN_400_b[i] <- mean(thr400$nNodesAG[tmp])
  SO_400_b[i] <- Mode(thr400$streamOrder[tmp])
  DD_400_b[i] <- mean(thr400$drainageDensity[tmp])
}

bin_thr500 <- seq(log10(min(thr500$thrValues)/max(thr500$thrValues)),0,length.out = nbins + 1)
thr_500_b <- nN_500_b <- SO_500_b <- DD_500_b <- numeric(nbins)
for (i in 1:nbins){
  if (i < nbins){
    tmp <- which(log10(thr500$thrValues/max(thr500$thrValues)) >= bin_thr500[i] & 
                   log10(thr500$thrValues/max(thr500$thrValues)) < bin_thr500[i+1])
  } else {
    tmp <- which(log10(thr500$thrValues/max(thr500$thrValues)) >= bin_thr500[i] & 
                   log10(thr500$thrValues/max(thr500$thrValues)) <= bin_thr500[i+1]) 
  }
  thr_500_b[i] <- mean(thr500$thrValues[tmp])
  nN_500_b[i] <- mean(thr500$nNodesAG[tmp])
  SO_500_b[i] <- Mode(thr500$streamOrder[tmp])
  DD_500_b[i] <- mean(thr500$drainageDensity[tmp])
}

bin_a250 <- seq(log10(min(a250)),log10(max(a250)),length.out = nbins+1)
a_250_b <- P_250_b <- numeric(nbins)
for (i in 1:nbins){
  if (i < nbins){
    tmp <- which(log10(a250) >= bin_a250[i] & log10(a250) < bin_a250[i+1])
  } else {
    tmp <- which(log10(a250) >= bin_a250[i] & log10(a250) <= bin_a250[i+1])  
  }
  a_250_b[i] <- mean(a250[tmp])
  P_250_b[i] <- mean(P250[tmp])
}

bin_a300 <- seq(log10(min(a300)),log10(max(a300)),length.out = nbins+1)
a_300_b <- P_300_b <- numeric(nbins)
for (i in 1:nbins){
  if (i < nbins){
    tmp <- which(log10(a300) >= bin_a300[i] & log10(a300) < bin_a300[i+1])
  } else {
    tmp <- which(log10(a300) >= bin_a300[i] & log10(a300) <= bin_a300[i+1])  
  }
  a_300_b[i] <- mean(a300[tmp])
  P_300_b[i] <- mean(P300[tmp])
}
bin_a400 <- seq(log10(min(a400)),log10(max(a400)),length.out = nbins+1)
a_400_b <- P_400_b <- numeric(nbins)
for (i in 1:nbins){
  if (i < nbins){
    tmp <- which(log10(a400) >= bin_a400[i] & log10(a400) < bin_a400[i+1])
  } else {
    tmp <- which(log10(a400) >= bin_a400[i] & log10(a400) <= bin_a400[i+1])  
  }
  a_400_b[i] <- mean(a400[tmp])
  P_400_b[i] <- mean(P400[tmp])
}
bin_a500 <- seq(log10(min(a500)),log10(max(a500)),length.out = nbins+1)
a_500_b <- P_500_b <- numeric(nbins)
for (i in 1:nbins){
  if (i < nbins){
    tmp <- which(log10(a500) >= bin_a500[i] & log10(a500) < bin_a500[i+1])
  } else {
    tmp <- which(log10(a500) >= bin_a500[i] & log10(a500) <= bin_a500[i+1])  
  }
  a_500_b[i] <- mean(a500[tmp])
  P_500_b[i] <- mean(P500[tmp])
}

### Plot figure
colors <- hcl.colors(4,palette="Zissou 1")
col250 <- colors[1]; col300 <- colors[2]; col400 <- colors[3]; col500 <- colors[4]
par(mfrow=c(2,2))

## Plot Number of nodes vs. threshold area
suppressWarnings(
  plot(thr_500_b/max(thr500$thrValues),nN_500_b,log="xy",pch=16,col=col500,cex=1,
       xlim=c(1e-6,1),ylim=c(1,1e6),bty="n",xaxt="n",yaxt="n",
       xlab="aT",ylab="Number of nodes at the AG level"))
suppressWarnings(
  points(thr_400_b/max(thr400$thrValues),nN_400_b,pch=16,col=col400,cex=0.8))
suppressWarnings(
  points(thr_300_b/max(thr300$thrValues),nN_300_b,pch=16,col=col300,cex=0.6))
suppressWarnings(
  points(thr_250_b/max(thr250$thrValues),nN_250_b,pch=16,col=col250,cex=0.4))
axis(1,pos=1)
axis(2,pos=1e-6)
lines(c(0.02,0.02),c(1,1e6),col="black")

ind250 <- which(thr250$thrValues/max(thr250$thrValues)<=0.02 & thr250$thrValues > thr250$thrValues[1])
ind300 <- which(thr300$thrValues/max(thr300$thrValues)<=0.02 & thr300$thrValues > thr300$thrValues[1])
ind400 <- which(thr400$thrValues/max(thr400$thrValues)<=0.02 & thr400$thrValues > thr400$thrValues[1])
ind500 <- which(thr500$thrValues/max(thr500$thrValues)<=0.02 & thr500$thrValues > thr500$thrValues[1])

all_aT <- c(thr250$thrValues[ind250]/max(thr250$thrValues), 
            thr300$thrValues[ind300]/max(thr300$thrValues),
            thr400$thrValues[ind400]/max(thr400$thrValues), 
            thr500$thrValues[ind500]/max(thr500$thrValues))
all_nN <- c(thr250$nNodesAG[ind250], thr300$nNodesAG[ind300], 
            thr400$nNodesAG[ind400], thr500$nNodesAG[ind500])

lmod <- lm(log(all_nN) ~ log(all_aT))
summary(lmod)
a_nN <- as.vector(lmod$coefficients)
lines(c(1e-6,0.565),exp(a_nN[1])*c(1e-6^a_nN[2],0.565^a_nN[2]),lty=2)
lines(c(0.02,0.02),c(0,10),col="black")
text(1e-5, 50, labels=sprintf("y = %.3f x + %.3f",a_nN[2],a_nN[1]))
text(1e-5, 10, labels=sprintf("R^2 = %.3f",summary(lmod)$r.squared))

## Plot Max stream order vs. threshold area
plot(thr_500_b/max(thr500$thrValues),SO_500_b,log="x",pch=16,col=col500,cex=1,
     xlim=c(1e-6,1),ylim=c(0,10),bty="n",xaxt="n",yaxt="n",
     xlab="aT",ylab="Maximum Stream Order")
points(thr_400_b/max(thr400$thrValues),SO_400_b,pch=16,col=col400,cex=0.8)
points(thr_300_b/max(thr300$thrValues),SO_300_b,pch=16,col=col300,cex=0.6)
points(thr_250_b/max(thr250$thrValues),SO_250_b,pch=16,col=col250,cex=0.4)
axis(1,pos=0)
axis(2,pos=1e-6)

all_SO <- c(thr250$streamOrder[ind250], thr300$streamOrder[ind300], 
            thr400$streamOrder[ind400], thr500$streamOrder[ind500])

lmod <- lm(all_SO ~ log(all_aT))
summary(lmod)
a_SO <- as.vector(lmod$coefficients)
lines(c(1e-6,1),a_SO[1]+a_SO[2]*log(c(1e-6,1)),lty=2)
lines(c(0.02,0.02),c(0,10),col="black")
text(1e-5, 2, labels=sprintf("y = %.3f x + %.3f",a_SO[2],a_SO[1]))
text(1e-5, 1, labels=sprintf("R^2 = %.3f",summary(lmod)$r.squared))


## Drainage density vs. Threshold area
plot(thr_500_b/thr500$thrValues[1],DD_500_b*sqrt(thr500$thrValues[1]),
     log="xy",pch=16,col=col500,xlim=c(1,1e6),ylim=c(1e-6,1.2),cex=1,bty="n",xaxt="n",yaxt="n",
     xlab="A_T [number of pixels]",ylab="Drainage density [pixel length^{-1}]")
points(thr_400_b/thr400$thrValues[1],DD_400_b*sqrt(thr400$thrValues[1]),pch=16,col=col400,cex=0.8)
points(thr_300_b/thr300$thrValues[1],DD_300_b*sqrt(thr300$thrValues[1]),pch=16,col=col300,cex=0.6)
points(thr_250_b/thr250$thrValues[1],DD_250_b*sqrt(thr250$thrValues[1]),pch=16,col=col250,cex=0.4)
lines(0.02*max(thr250$thrValues)/thr250$thrValues[1]*c(1,1),c(1e-6,1),col=col250)
lines(0.02*max(thr300$thrValues)/thr300$thrValues[1]*c(1,1),c(1e-6,1),col=col300)
lines(0.02*max(thr400$thrValues)/thr400$thrValues[1]*c(1,1),c(1e-6,1),col=col400)
lines(0.02*max(thr500$thrValues)/thr500$thrValues[1]*c(1,1),c(1e-6,1),col=col500)
axis(1,pos=1e-6)
axis(2,pos=1)

all_AT <- c(thr250$thrValues[ind250]/thr250$thrValues[1], thr300$thrValues[ind300]/thr300$thrValues[1],
            thr400$thrValues[ind400]/thr400$thrValues[1], thr500$thrValues[ind500]/thr500$thrValues[1])
all_DD <- c(thr250$drainageDensity[ind250]*sqrt(thr250$thrValues[1]), 
            thr300$drainageDensity[ind300]*sqrt(thr300$thrValues[1]), 
            thr400$drainageDensity[ind400]*sqrt(thr400$thrValues[1]), 
            thr500$drainageDensity[ind500]*sqrt(thr500$thrValues[1]))

lmod <- lm(log(all_DD) ~ log(all_AT))
summary(lmod)
a_DD <- as.vector(lmod$coefficients)
lines(c(1,1e6),exp(a_DD[1])*(c(1^a_DD[2],1e6^a_DD[2])),lty=2)
text(10, 1e-4, labels=sprintf("y = %.3f x + %.3f",a_DD[2],a_DD[1]))
text(10, 1e-5, labels=sprintf("R^2 = %.3f",summary(lmod)$r.squared))

## Scaling of areas
plot(a_500_b/a500[1],P_500_b,log="xy",pch=16,col=col500,cex=1,xlim=c(1,1e6),ylim=c(1e-6,1),
     bty="n",xaxt="n",yaxt="n",xlab="a [no. pixels]",ylab="P [A >= a]")
axis(1,pos=1e-6)
axis(2,pos=1)
points(a_400_b/a400[1],P_400_b,pch=16,col=col400,cex=0.8)
points(a_300_b/a300[1],P_300_b,pch=16,col=col300,cex=0.6)
points(a_250_b/a250[1],P_250_b,pch=16,col=col250,cex=0.4)
lines(0.02*max(a250)/a250[1]*c(1,1),c(1e-6,1),col=col250)
lines(0.02*max(a300)/a300[1]*c(1,1),c(1e-6,1),col=col300)
lines(0.02*max(a400)/a400[1]*c(1,1),c(1e-6,1),col=col400)
lines(0.02*max(a500)/a500[1]*c(1,1),c(1e-6,1),col=col500)
ind250 <- which(a250<=0.02*max(a250))
ind300 <- which(a300<=0.02*max(a300))
ind400 <- which(a400<=0.02*max(a400))
ind500 <- which(a500<=0.02*max(a500))
a_all <- c(a250[ind250]/a250[1], a300[ind300]/a300[1], a400[ind400]/a400[1], a500[ind500]/a500[1])
P_all <- c(P250[ind250], P300[ind300], P400[ind400], P500[ind500])
lmod <- lm(log(P_all) ~ log(a_all) )
summary(lmod)
B <- lmod$coefficients
lines(c(1,1e6),exp(B[1])*c(1^B[2],1e6^B[2]),lty=2)
text(10, 1e-4, labels=sprintf("y = %.3f x + %.3f",B[2],B[1]))
text(10, 1e-5, labels=sprintf("R^2 = %.3f",summary(lmod)$r.squared))

par(old.par)

# # # # # # # # # # # # # # # # # # # # # # # 
#### FIGURE 6 - CONVERSION TO IGRAPH, SSN ####
# # # # # # # # # # # # # # # # # # # # # # # 

cat("Creating Figure 6...\n")

set.seed(1)
OCN <- create_OCN(20, 20, outletPos = 3, cellsize = 500)
OCN <- landscape_OCN(OCN, slope0 = 0.01)
OCN <- aggregate_OCN(OCN, thrA = 5*500^2)

g <- OCN_to_igraph(OCN, level = "AG")
ssnOCN <- OCN_to_SSN(OCN, level = "RN", obsDesign = SSN::binomialDesign(50),
                     path = paste(tempdir(), "/",as.numeric(Sys.time()),".ssn", sep = ""), importToR = TRUE)

# Plot figure (SSN)
plot.SpatialStreamNetwork(ssnOCN, "upDist", breaktype = "user", brks = seq(0,15000,3000), 
                          asp = 1, bty="n", xaxt="n", yaxt="n", xlab="", ylab="") 
title("SSN object")
text(0.87,0.65,labels="Distance from outlet [m]")

## Plot figure (igraph)
par(mfrow=c(1,2), mai=c(0.1,0.1,0.1,0.1))
plot.igraph(g, vertex.color = hcl.colors(OCN$AG$nNodes, palette="Set 2"), 
            layout = matrix(c(OCN$AG$X,OCN$AG$Y),ncol = 2, nrow = OCN$AG$nNodes))
title("igraph object")

draw_thematic_OCN(c(1:OCN$AG$nNodes), OCN, discreteLevels = TRUE, drawNodes = TRUE,
                  colPalette = hcl.colors(OCN$AG$nNodes, palette="Set 2"),  cex = 4, riverColor = "#999999",
                  backgroundColor = NULL, addLegend = FALSE)
title("OCN object")
text(OCN$AG$X, OCN$AG$Y)

par(old.par)

# # # # # # # # # # # # # # # # # # # # #
#### FIGURE 7 - METAPOPULATION MODEL ####
# # # # # # # # # # # # # # # # # # # # # 

cat("Creating Figure 7...\n")

## Build OCN
set.seed(1)
OCN <- create_OCN(20, 20, outletPos = 3, cellsize = 500)
OCN <- landscape_OCN(OCN, slope0 = 0.01)
OCN <- aggregate_OCN(OCN, thrA = 5*500^2)
OCN <- paths_OCN(OCN, pathsRN = TRUE)
OCN <- rivergeometry_OCN(OCN, widthMax = 5) 

farthestNode <- which(OCN$RN$downstreamPathLength[ , OCN$RN$outlet] ==
                        max(OCN$RN$downstreamPathLength[ , OCN$RN$outlet]))[1]
dist_from_NW <- sqrt((OCN$RN$X-250)^2 + (OCN$RN$Y-20*500-250)^2)
otherHeadwater <- which(dist_from_NW == min(dist_from_NW))

## Weights for upstream movement
Y <- rep(1,OCN$RN$nNodes)                    
for (i in 1:OCN$RN$nNodes){
  if (i != OCN$RN$outlet){
    Y[i] <- OCN$RN$A[i]/(OCN$RN$W[ , OCN$RN$downNode[i]] %*% OCN$RN$A)
  }
}

## input data
K <- 10*OCN$RN$width                    # calculate carrying capacity 
pop0 <- numeric(OCN$RN$nNodes)          # initial random population vector
pop0[farthestNode] <- 1
nTimestep <- 1000                       # number of timesteps
r <- 1.05                               # proliferation rate
pd <- 0.5                               # probability to move downstream
pu <- 1 - pd                            # probability to move upstream
g <- 0.1                                #fraction of individuals moving


## metapopulation model
metapop_model <- function(pop0, r, K, g, pd, Y, OCN, nTimestep){
  pu <- 1 - pd
  pop <- matrix(data=0,ncol=nTimestep,nrow=OCN$RN$nNodes)
  pop[,1] <- pop0                                          
  for (t in 2:nTimestep){
    for (i in 1:OCN$RN$nNodes){
      pop[i, t] <-  r*pop[i, t-1]/(1 + pop[i, t-1]*(r-1)/K[i]) +
        - (pu*(sum(OCN$RN$W[ , i])>0) + pd*(sum(OCN$RN$W[i, ])>0)) * g * pop[i,t-1] +
        + pd * OCN$RN$W[ , i] %*% (g * pop[ , t-1]) +
        + pu * Y[i] * OCN$RN$W[i, ] %*% (g * pop[ , t-1])
    }
  }
  return(pop)
}

## run model
pop1 <- metapop_model(pop0, r, K, g, pd, Y, OCN, nTimestep=800)
pop2 <- metapop_model(pop0, r, K, g, pd=0.7, Y, OCN, nTimestep=800)

## evaluate equilibrium population sizes
conv_pop1_red <- which(pop1[OCN$RN$outlet,]>0.99*pop1[OCN$RN$outlet,800])[1]
conv_pop2_red <- which(pop2[OCN$RN$outlet,]>0.99*pop2[OCN$RN$outlet,800])[1]
conv_pop1_green <- which(pop1[farthestNode,]>0.99*pop1[farthestNode,800])[1]
conv_pop2_green <- which(pop2[farthestNode,]>0.99*pop2[farthestNode,800])[1]
conv_pop1_blue <- which(pop1[otherHeadwater,]>0.99*pop1[otherHeadwater,800])[1]
conv_pop2_blue <- which(pop2[otherHeadwater,]>0.99*pop2[otherHeadwater,800])[1]
conv_metapop1 <- which(colSums(pop1)>0.99*sum(pop1[,800]))[1]
conv_metapop2 <- which(colSums(pop2)>0.99*sum(pop2[,800]))[1]

### Plot figure (time evolution)
colors <- hcl.colors(3,palette="Set 2")
par(mfrow = c(2, 1), mai = c(1, 1, 1,1))

plot(pop1[OCN$RN$outlet, ], type = "l", ylim = c(0, 80), col = colors[1], 
     xlab = "Time", ylab = "Population", lwd = 2, bty="n",xaxt="n",yaxt="n")
title("Evolution of local population size")
axis(1,pos=0); axis(2,pos=0)
lines(c(100,100),c(0,80),lwd=0.5)
lines(c(250,250),c(0,80),lwd=0.5)
lines(c(600,600),c(0,80),lwd=0.5)

lines(c(conv_pop1_red,conv_pop1_red),c(0,80),lwd=0.5,col=colors[1])
lines(c(conv_pop2_red,conv_pop2_red),c(0,80),lwd=0.5,col=colors[1], lty = 2)
lines(c(conv_pop1_green,conv_pop1_green),c(0,80),lwd=0.5,col=colors[2])
lines(c(conv_pop2_green,conv_pop2_green),c(0,80),lwd=0.5,col=colors[2], lty = 2)
lines(c(conv_pop1_blue,conv_pop1_blue),c(0,80),lwd=0.5,col=colors[3])
lines(c(conv_pop2_blue,conv_pop2_blue),c(0,80),lwd=0.5,col=colors[3], lty = 2)

lines(pop2[OCN$RN$outlet, ], type = "l", col = colors[1],  lwd = 2, lty = 2)
lines(pop1[farthestNode, ], type="l", col=colors[2],lwd=2)
lines(pop2[farthestNode, ], type="l", col=colors[2],lwd=2, lty = 2)
lines(pop1[otherHeadwater, ], type="l", col=colors[3],lwd=2)
lines(pop2[otherHeadwater, ], type="l", col=colors[3],lwd=2, lty = 2)

plot(colSums(pop1), type = "l", xlab = "Time", ylab = "Population", lwd = 2, 
     ylim = c(0, 1500), bty="n",xaxt="n",yaxt="n",col="#ACA2EC")
lines(colSums(pop2), lty = 2, type = "l", lwd = 2,col="#ACA2EC")
title("Evolution of metapopulation size")
axis(1,pos=0); axis(2,pos=0)
lines(c(100,100),c(0,1500),lwd=0.5)
lines(c(250,250),c(0,1500),lwd=0.5)
lines(c(600,600),c(0,1500),lwd=0.5)
lines(c(conv_metapop1,conv_metapop1),c(0,1500),lwd=0.5,col="#ACA2EC")
lines(c(conv_metapop2,conv_metapop2),c(0,1500),lwd=0.5,col="#ACA2EC",lty=2)

### Plot figure (spatial spread)
par(mfrow = c(3, 2), mai = c(0.1, 0, 0.2, 0))

draw_thematic_OCN(pop1[,100], OCN, colLevels = c(0, 80, 1000),
                  drawNodes = TRUE, backgroundColor = NULL)
points(OCN$RN$X[OCN$RN$outlet],OCN$RN$Y[OCN$RN$outlet],pch=1,col=colors[1],cex=2.5,lwd=3,asp=1)
points(OCN$RN$X[farthestNode],OCN$RN$Y[farthestNode],pch=1,col=colors[2],cex=2.5,lwd=3,asp=1)
points(OCN$RN$X[otherHeadwater],OCN$RN$Y[otherHeadwater],pch=1,col=colors[3],cex=2.5,lwd=3,asp=1)
title("Time = 100")
draw_thematic_OCN(pop2[,100], OCN, colLevels = c(0, 80, 1000),
                  drawNodes = TRUE, backgroundColor = NULL)
title("Time = 100")

draw_thematic_OCN(pop1[,250], OCN, colLevels = c(0, 80, 1000),
                  drawNodes = TRUE, backgroundColor = NULL)
title("Time = 250")
draw_thematic_OCN(pop2[,250], OCN, colLevels = c(0, 80, 1000),
                  drawNodes = TRUE, backgroundColor = NULL)
title("Time = 250")

draw_thematic_OCN(pop1[,600], OCN, colLevels = c(0, 80, 1000),
                  drawNodes = TRUE, backgroundColor = NULL)
title("Time = 600")
draw_thematic_OCN(pop2[,600], OCN, colLevels = c(0, 80, 1000),
                  drawNodes = TRUE, backgroundColor = NULL)
title("Time = 600")

par(old.par)
```
